# Large-scale docking predicts that sORF-encoded peptides may function through protein-peptide interactions in Arabidopsis thaliana

**DOI:** 10.1101/335687

**Authors:** Rashmi R. Hazarika, Nikolina Sostaric, Yifeng Sun, Vera van Noort

**Affiliations:** KU Leuven, Department of Microbial and Molecular Systems, KU Leuven Kasteelpark Arenberg 22, B-3001 Leuven, Belgium.; KU Leuven, Faculty of Engineering Technology, Campus Group T, Andreas Vesaliusstraat 13, 3000 Leuven, Belgium.; Leiden University, Institute of Biology Leiden, 2300 RA Leiden, The Netherlands.

## Abstract

Several recent studies indicate that small Open Reading Frames (sORFs) embedded within multiple eukaryotic non-coding RNAs can be translated into bioactive peptides of up to 100 amino acids in size. However, the functional roles of the 607 Stress Induced Peptides (SIPs) previously identified from 189 Transcriptionally Active Regions (TARs) in *Arabidopsis thaliana* remain unclear. To provide a starting point for function annotation of these peptides, we performed a large-scale prediction of peptide binding sites on protein surfaces using and coarse-grained peptide docking. The docked models were subjected to further atomistic refinement and binding energy calculations. A total of 530 peptide-protein pairs were successfully docked. In cases where a peptide encoded by a TAR is predicted to bind at a known ligand or cofactor-binding site within the protein, it can be assumed that the peptide modulates the ligand or cofactor-binding. Moreover, we predict that several peptides bind at protein-protein interfaces, which could therefore regulate the formation of the respective complexes. Protein-peptide binding analysis further revealed that peptides employ both their backbone and side chain atoms when binding to the protein, forming predominantly hydrophobic interactions and hydrogen bonds. In this study, we have generated novel predictions on the potential protein-peptide interactions in *A. thaliana*, which will help in further experimental validation.

**Author summary:** Due to their small size, short peptides are difficult to find and have been ignored in genome annotations. Only recently, we have realized that these short peptides of less than 100 amino acids may actually play an important role in the cell. Currently, there are no high-throughput methods to find out what the functions of these peptides are in contrast with efforts that exist for ‘normal’proteins. In this work, we try to fill this gap by predicting with which larger proteins, the short peptides might interact to exert their function. We find that many peptides bind to pockets where normally other proteins or molecules bind. We thus think that these peptides that are induced by stress, may regulate protein-protein and protein-molecule binding. We make this information available through our database ARA-PEPs so that individual predictions can be followed up.

## Introduction

Over the years, the functional importance of short plant signaling peptides has been overshadowed by other groups of molecules. For instance, the phytohormone auxin was shown to be involved in bidirectional polar transport across tissues, controlling plant growth-related processes (Grunewald & Friml, 2010; Murphy *et al*, 2012). Furthermore, microRNAs are considered to be important signaling molecules, regulating developmental processes in plants by moving from one cell to another over long distances (Marín-González & Suárez-López, 2012). It was only within the last decade that the roles of plant peptides in a wide variety of cellular functions were established by multiple studies (Matsubayashi, 2011; Tavormina *et al*, 2015). Some of these peptides may be encoded by short Open Reading Frames (sORFs), that were earlier assumed to be non-coding (Amor *et al*, 2009; Chen *et al*, 2015; Ladoukakis *et al*, 2011; Crappé *et al*, 2013; Ruiz-Orera & Messeguer, 2014; Andrews & Rothnagel, 2014). Several recent studies clearly demonstrate that sORFs embedded within non-coding RNAs (ncRNAs), intergenic regions and pseudogenes can indeed be translated into bioactive peptides. In our previous work, we have identified several Transcriptionally Active Regions (TARs) induced upon the application of biotic (*Botrytis cinerea*) and abiotic stress (Paraquat) in *Arabidopsis thaliana*. These TARs could be translated into Stress-Induced Peptides (SIPs), which can be specifically categorized depending on the applied stress condition into *Botrytis cinerea* Induced Peptides (BIPs) and Oxidative Stress Induced Peptides (OSIPs) that we catalogued in a database ARA-PEPs (Hazarika *et al*, 2017; De Coninck *et al*, 2013). Although some physiological effects of sORF-encoded peptides have been discovered, the molecular mechanism by which they exert their function through interaction with other molecules is largely unknown. We postulate that the peptides could work through interactions with proteins, as protein-peptide interactions have previously been well established as important mediators of protein-protein interactions, partaking in signal transduction, cell-to-cell communication, protein trafficking and other regulatory pathways (London *et al*, 2010; Schindler *et al*, 2015; Kilburg & Gallicchio, 2016; Petsalaki *et al*, 2009; Neduva & Russell, 2005; Perkins *et al*, 2010; Pawson & Nash, 2003).

Peptide-mediated interactions constitute 15-40% of all protein-protein interactions (Petsalaki and Russell 2008, Neduva and Russell, 2005). Most of the studies performed so far in order to understand protein-peptide interactions focused on small peptides that may be short linear recognition motifs originating from disordered protein regions (Kilburg & Gallicchio, 2016; London *et al*, 2010). Investigating those interactions is experimentally challenging, and this has led to limited progress in the field of protein-peptide interactions validation. On the other side, the successful modeling of such complexes depends on prior structural knowledge of the protein that acts as a receptor. A number of protein-peptide docking methods, such as Rosetta FlexPepDock (Raveh *et al*, 2010, 2011), GalaxyPepDock (Ko *et al*, 2012; Heo *et al*, 2013; Lee *et al*, 2015), MedusaDock (Ding *et al*, 2010), DynaDock (Antes, 2010), CABS-dock (Kurcinski *et al*, 2015; Wabik *et al*, 2015), pepATTRACT (Schindler *et al*, 2015), HADDOCK (Dominguez *et al*, 2003; Trellet *et al*, 2013), and tools to predict binding sites on proteins, such as PepSite2 (Trabuco *et al*, 2012; Petsalaki *et al*, 2009), have been developed. Moreover, curated data also exists for characterization of protein-peptide interactions, e.g. a non-redundant database of high-resolution peptide-protein complexes called the *peptiDB* (London *et al*, 2010). Although docking strategies are the preferred methods for predicting protein-peptide interactions, they are associated with certain limitations, such as difficulty in docking peptides longer than 4 amino acids, owing to their high degree of conformational flexibility. In our current study, we opted to combine the peptide-protein docking method pepATTRACT-local with binding site predictions obtained from the PepSite2 server,that uses training data of known protein-peptide complexes from Protein Data Bank (PDB) to define Spatial Position Specific Scoring Matrices (S-PSSMs). Furthermore, as biological systems are not static, we also looked into dynamics of the obtained docked models and calculated the energetics of binding based on multiple conformations that the protein-peptide system can acquire in the solution.

We hypothesize that a SIP encoded by a TAR may bind on a protein at one of its pockets, or to a known ligand or cofactor-binding site, and consequently affect the function of the protein as a whole. Moreover, peptides may bind at the interfaces of multi-chain complexes and modulate their activity. Protein-peptide interactions involve smaller interfaces whose affinity is usually weaker and are transient as they can rapidly make and break interactions in response to sudden cellular perturbations, for instance stress conditions (Stein & Aloy, 2008; Perkins *et al*, 2010).

In recent years, there has been a growing interest in developing protein-protein interaction inhibitors based on peptides or peptide derivatives. Molecules that can mimic the binding or functional sites of proteins are promising candidates for different types of biological applications. Synthetic peptides are widely choiced molecules for mimicry of protein sites because they can be easily synthesized as exact copies of protein fragments, or they may be generated by introducing diverse chemical modifications to the peptide sequence, and/or by modifying the peptide backbone (Groß *et al*, 2016). Peptide mimics have the potential to be developed as attractive targets for agriculture, especially plant disease control, and for therapeutic interventions (Beekman & Howell, 2016).

As detailed above, when no information about the peptide-binding site on protein receptors is available, there is need for computational approaches to predict peptide-binding sites on protein surfaces, as these models can serve as starting points for experimental characterization of novel protein-peptide interactions. This will be especially beneficial in studying the model plant *A. thaliana*, in which peptides have multiple important roles, but have been understudied till now. Molecular docking studies can be used effectively to explore the binding mode of putative peptides onto proteins, serving as an excellent approach for *de novo* design of peptides targeting various other biosynthetic pathways in major eukaryotes. In the current study, we investigate the potential roles of sORF-encoded stress induced peptides in targeting the key regulatory enzymes, as this could further indicate their roles in mediating the stress-response mechanisms.

## Results

### Short peptides may exert their function by interacting with proteins

In a previous study, we identified 189 TARs in response to plant oxidative stress by the herbicide Paraquat and the fungus *Botrytis cinerea*, which could be translated into 607 SIPs (Hazarika *et al*, 2017; De Coninck *et al*, 2013). A peptide fragment library consisting of 23,113 k-mers, ranging from 4 to 10 amino acid residues, was generated and searched for potential binding sites on *A. thaliana* proteins from the PDB repository, using PepSite2. We screened 12,540,140 protein-peptide pairs and found 3,769,393 significant matches at PepSite2 score > 60 and *p*-value <= 0.1. We additionally screened for short peptide motif matches on *A. thaliana* proteins using BLASTP and found 302 matches. We performed initial docking analysis using the pepATTRACT protocol, and a larger subset of 576 protein-peptide pairs was devised by pooling together the above 302 protein-peptide pairs as well as others with significant PepSite2 score (Figure 1A). The list of docked pairs can be accessed through the url (https://www.biw.kuleuven.be/CSB/ARA-PEPs/SIP_PDB_interactions.php).In our study, 46 protein-peptide complexes failed to dock, and the reason could be the large conformational changes of the protein caused by binding of a flexible peptide; this indeed remains a big problem of docking methods (Trellet *et al*, 2013). From the docked models, we filtered out the 104 top protein-peptide pairs, and for each of them characterized the protein-peptide binding, and determined the free energy of binding. Our results show that there exists a huge repertoire of potential peptide binding sites on all available *A. thaliana* proteins out of which we explored only 0.015%. Large-scale predictions of potential protein-peptide pairs can aid in future experimental validations for understanding cell-to-cell communication during plant development or stress-tolerance mechanisms.

**Figure 1.**
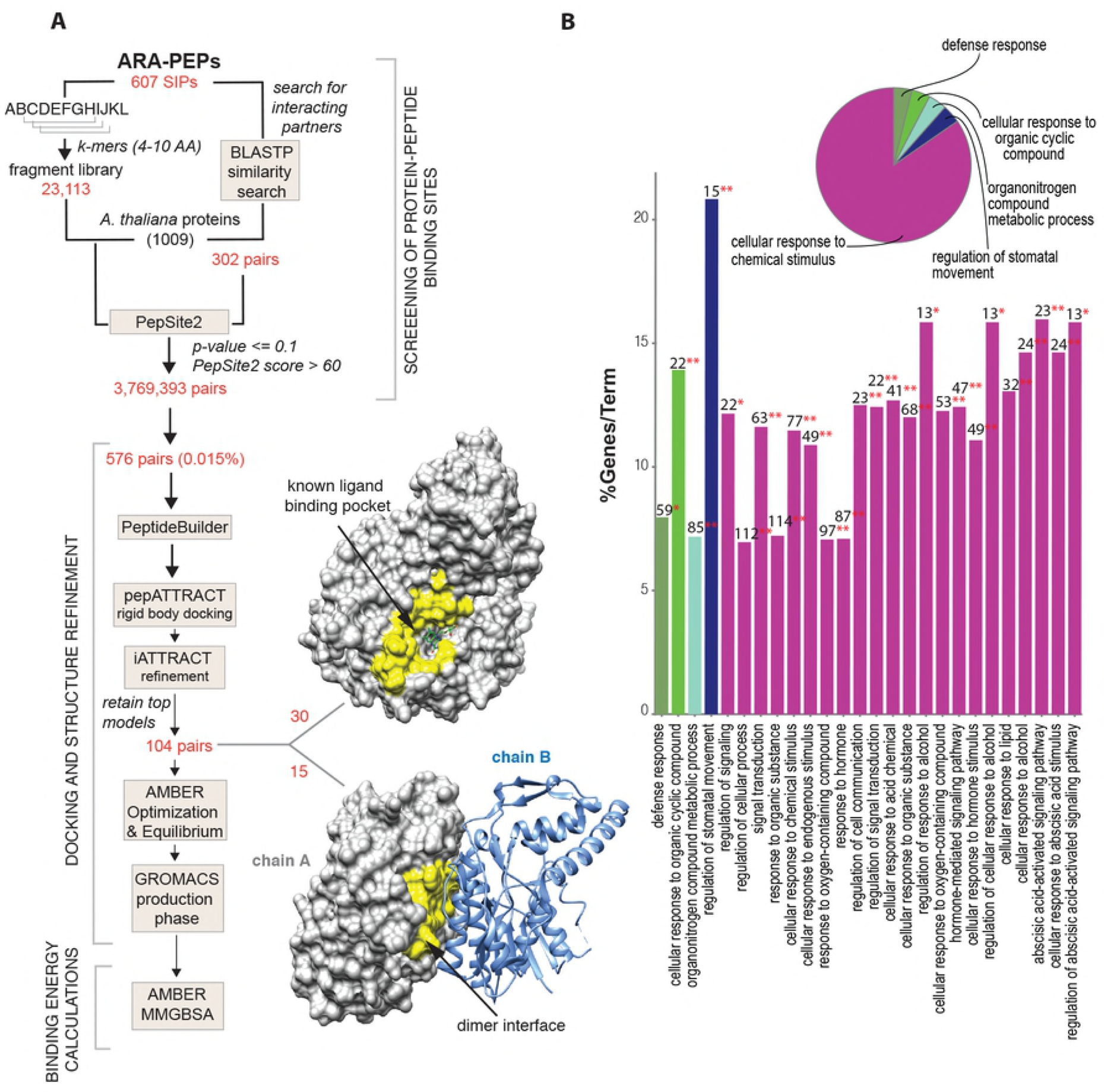
**(A)** Computational pipeline for prediction of protein-peptide pairs. 23,113 k-mers were screened for binding sites on 1009 *A. thaliana* proteins. Out of 3,769,393 significant matches, 576 pairs were docked, and 104 pairs were further studied in detail. 30 peptides may bind to a ligand binding/catalytic site on a protein and 15 peptides may bind at the dimer interface between 2 chains of a protein complex. The peptide binding pocket is highlighted in yellow. **(B)** Histogram showing specific GO terms related to the associated proteins from protein-peptide screening analysis. The bars represent the number of proteins from the analyzed cluster associated with the term, and the label displayed on the bars is the percentage of proteins compared to all proteins associated with the term. The overview pie-chart presents functional groups for the proteins where the name of the group is given by the most significant term in the group. GO enrichment analysis revealed 5 main groups and each group section in the pie-chart correlates with the number of terms in each group.

Specific inhibitors may mimic portions of protein interfaces and can bind to a peptide binding pocket located at the interface between two monomers. In our study we found that 15 peptides bind at the interface between subunits of protein complexes, indicating the ability to modulate complex’ s activity (Figure 1A, Supplementary table 2). Among the 15 models, in three of the cases the peptides bind in a similar way to known characterized short peptides or portion of a full-length protein (Supplementary Figure 3). We also observed that 30 peptides bind to a known ligand/cofactor binding site (Figure 1A, Supplementary table 1). Ligand and protein binding sites may often overlap within protein families as it has been shown that a peptide may compete with the ligand for the binding site or non-competitively bind to the pocket along with the ligand molecule. The structural models obtained in the current study will facilitate future validations as we can directly compare the binding modes of peptides on proteins. While we observe that both pepATTRACT-local and pepATTRACT-blind produce similar results for 56% of the docked pairs, the other 44% of the pairs were not docked at the same site of the protein using the two methods. Under assumption that the PepSite2 correctly indicated the binding regions, these results show how the use of restraints in the docking protocol can help in concentrating the search around relevant regions of the protein-peptide interaction space. Other reports have previously shown that restraint-based dockings yield better results as compared to blind docking methods (Vazda and Kozakov, 2009, de Vries et al., 2007).

**Figure 2.**
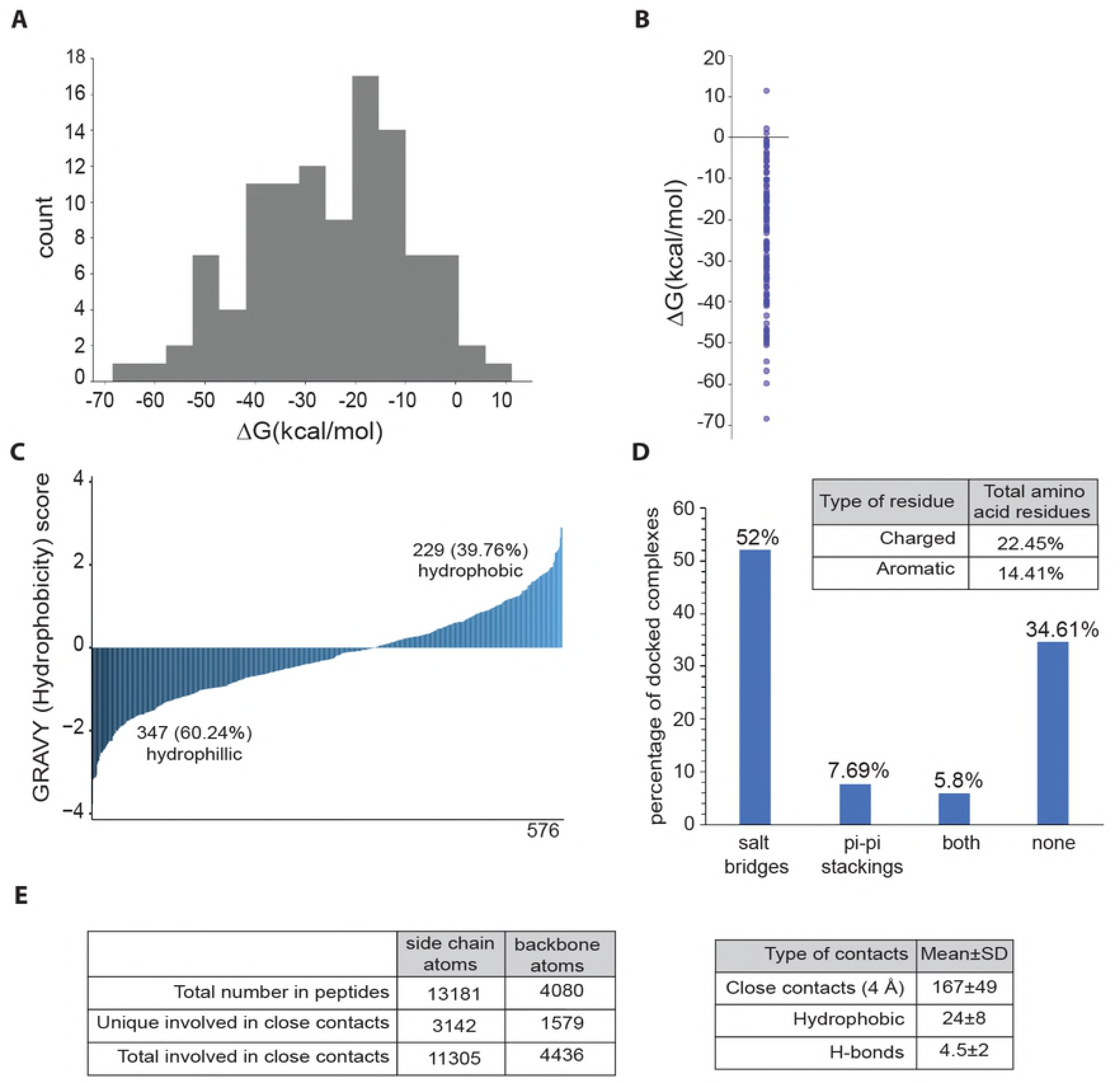
Characterization of peptides and protein-peptide binding interactions. **(A)** Histogram showing Δ*G*_bind_ values for 104 top models **(B)** Δ*G*_bind_ values as individual data points **(C)** Hydropathy index for all the 576 peptides predicted to interact with *A. thaliana* proteins **(D)** Total number of charged and aromatic residues in the peptides that interact with proteins. The plot also shows the total number of salt-bridges and pi-pi stackings formed in the top models. **(E)** Different types of interactions formed by the protein-peptide pairs

**Figure 3.**
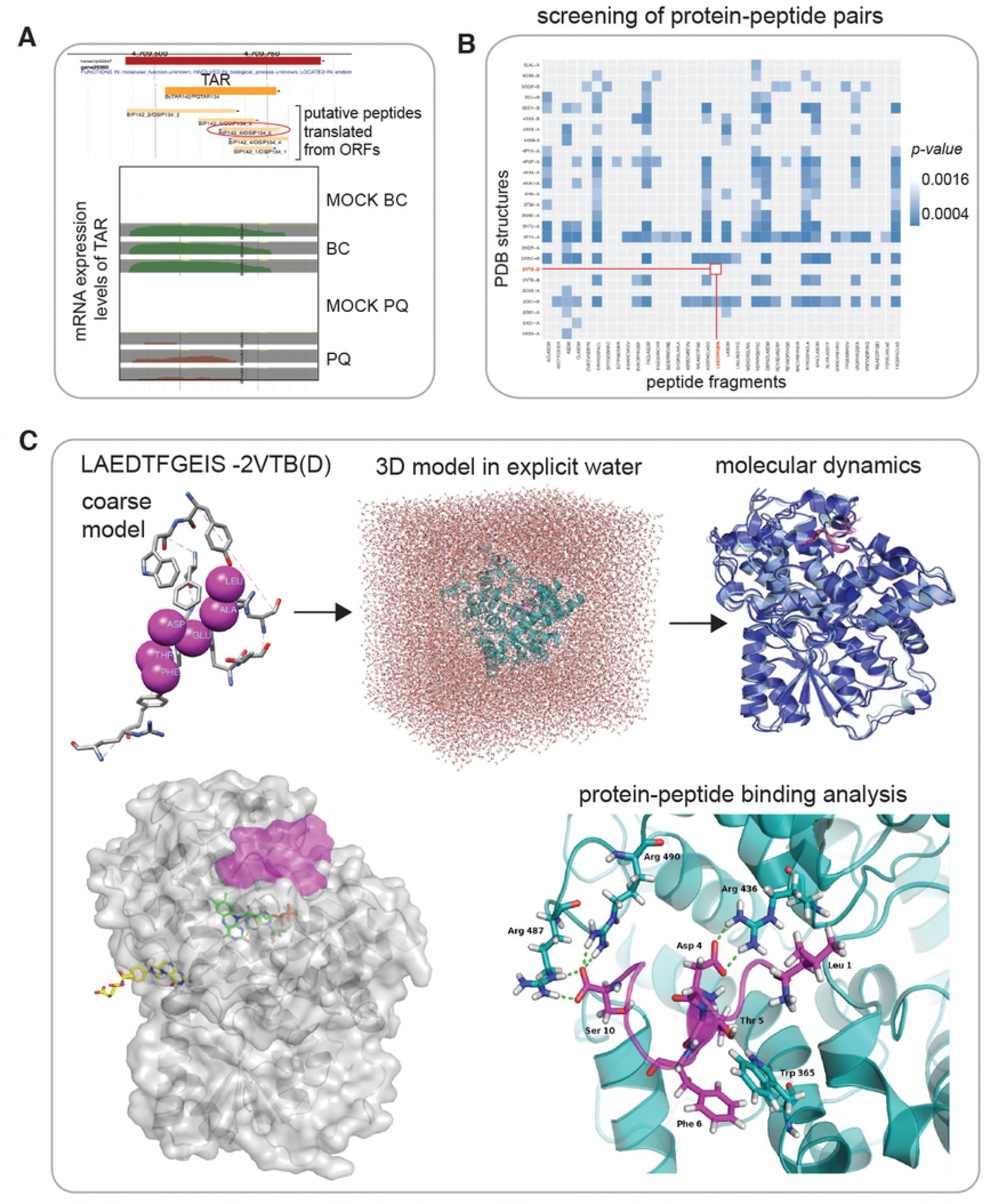
**(A)** Overview of Transcriptionally Active Region BcTAR142/PQTAR134 induced under stress conditions which might encode a short peptide. The TAR shows mRNA expression levels under treated (PQ and BC) and mock conditions (mock_BC and mock_PQ). **(B)** Screening of all possible peptide fragments from BIP142_3/OSIP134_3 against all *A. thaliana* proteins in PDB. **(C)** A coarse model of peptide LAEDTFGEIS bound to CRYD protein (chain D of 2VTB). Restraint based docking was performed, followed by surrounding of the 3D model with explicit water. The solvated structure was optimized and then used for MD simulation. The conformational snapshots from the MD were used to calculate Δ*G*_bind_ value for protein-peptide binding, and for visual inspection of the mode of binding (details in the text). Finally, superposition of the docked model (CRYD as grey, and docked LAEDTFGEIS as magenta surface) and the original PDB structure, which has FAD (green) and MTHF (yellow) co-factors bound, suggests that peptide might have an effect on FAD binding, and consequently on CRYD’s function.

In addition, we investigated if peptide binding pockets on proteins could bind multiple peptides, or in other words, whether a peptide prefers to bind to one specific pocket on a protein. To test this, we generated a randomly shuffled list of peptides while keeping the list of PDB structures intact, followed by scanning for binding sites using PepSite2. In 95.15% of the cases the peptides preferred to bind to the same pocket, while in only 4.84% of the cases the peptides were bound to different pockets on the same receptor (Supplementary Figure 1A). It is possible that the S-PSSMs capture the binding modes of amino acids in such a way that amino acids in a peptide sequence may prefer to bind to chemically similar binding sites on proteins, e.g. hydrophobic amino acids from the peptide tend to bind hydrophobic protein regions (Petsalaki *et al*, 2009) of appropriate sizes. While some reports suggest that peptides often look for a large enough pocket to bind followed by latching onto it with the help of a few hotspot residues (London et al, 2010, 2012), other reports suggest that several different peptides are able to bind to the same protein domain by exhibiting special properties such as promiscuity (Bhattacherjee & Wallin, 2013). Furthermore, the seemingly more important role of peptide backbone compared to side chain atoms (detailed below) in protein binding provides another explanation for the observed promiscuity, as backbone atoms are the same independent of the amino acid sequence of the peptide.

We mapped 835 unique *A. thaliana* protein chains from the initial screening to Uniprot IDs using annotations from SIFTS database (Velankar *et al*, 2013) and performed Gene Ontology (GO) enrichment analysis using REACTOME_Pathways and GO_BiologicalProcess ontology. GO analysis of the *A. thaliana* proteins with significant scores revealed that they may be categorized into 5 main groups *viz.* defense response, cellular response to organic cyclic compound, organonitrogen compound metabolic process, regulation of stomatal movement and cellular response to chemical stimulus (Figure 1B).

### Characterization of protein-peptide binding interactions

We carried out a general characterization of protein-peptide interactions in the 104 docked pairs. We extracted top 10 models from the docking results after iATTRACT refinement and calculated the average number of interactions within each docked pair. Each receptor atom that comes within 4 Å of any ligand atom is considered as a close contact. We determined the mean number of close contacts per docked pair to be 167±49. All protein-peptide pairs interacted with each other using hydrogen bonds and hydrophobic interactions. The mean number of hydrophobic interactions per system is 24±8, and the mean number of hydrogen bonds per docked pair is 4.5±2, where donors of hydrogen bonds are localized on protein in 51% of the cases (Figure 2E). While hydrogen bonds and hydrophobic interactions are omnipresent, not all pairs formed salt bridges and pi-pi stackings (Figure 2E and 2D). Within the peptides in our top 104 models, 22.5% of the total amino acid residues are charged (Arg, Lys, His, Asp, Glu) and 14.4% residues are aromatic (His, Phe, Tyr, Trp). In agreement with the amino acid composition, we also found salt bridges to be more prevalent than pi-pi stackings in the docked systems: 52% of protein-peptide pairs contain salt bridges with the mean number per system 1.72±1, and 7.7% form pi-pi stackings with the mean of 1.63±0.7. An additional 5.8% of pairs have both salt bridges and pi-pi stackings, while the remaining 34.6% do not form any salt bridges or pi-pi stackings (Figure 2D).

In total, 23.8% of peptide side chain and 38.7% of backbone atoms participate in close contacts, with an average number of contacts per interacting atom being 3.6 and 2.8, respectively (Figure 2E). In average, the ratio of the unique peptide side chain:backbone atoms involved in close contacts is 2:1 for the top model in 104 docked systems, while the overall side chain:backbone ratio of atoms is 3.2:1. If side chain and backbone atoms of the peptide were equally important in protein binding, we would expect the ratio of atoms involved in close contacts to also be 3.2:1. Instead, its lower value suggests that peptide backbone atoms might be more important in protein-peptide interactions than the side chain ones.

We determined the overall hydropathy index for all the peptide fragments (576) and found that 60.2% of the peptides are hydrophilic, while the remaining 39.8% are hydrophobic in nature (Figure 2C). A larger fraction of SIPs in our dataset have high hydrophilicity or a lower GRAVY index score, suggesting that they may mainly interact with globular proteins rather than with hydrophobic regions that spans membranes. Peptides with fewer ionic/charged groups are generally less soluble in water and are therefore prone to aggregation and interacting with hydrophobic pockets of larger proteins.

After analysis of the docked models, we took a further look into the dynamics of the top model from each of the 104 protein-peptide docked pairs and calculated the free energy of binding based on 100 conformational snapshots from molecular dynamics for each system (Figure 2A, B). Per-residue decomposition of protein-peptide binding energies also allowed identification of amino acid types that frequently (in multiple systems) have significant binding contribution (Supplementary Figure 2A). For instance, arginine residue, located in proteins at the peptide binding interface, stands out as a recurring amino acid with significant stabilizing effect on the binding (negative value of the Δ*G*_bind_ contribution). Interestingly, a prevalent contribution of negatively charged peptide amino acids is seemingly lacking. Visual investigation of trajectories obtained by molecular dynamics shows that arginines make salt bridges with negatively charged carboxyl groups of peptide C-terminal amino acids in 70% of cases (31/44), making this interaction independent of amino acid type present in the peptide. Other prominent protein residues that predominantly stabilize interactions with the peptides are the charged (Glu, Asp) and aromatic ones (Trp, Phe, Tyr).

Local destabilizing effect on binding is shown by different peptide amino acids, containing side chains of largely different properties (Supplementary Figure 2A). However, a more detailed view reveals that this is a consequence of amino acid location within a peptide, rather than its chemical composition (Supplementary Figure 2B). In average, non-terminally located amino acids contribute to the binding in a stabilizing manner, N-termini destabilize protein-peptide interaction, while C-terminal amino acids have different average effect depending on amino acid type, and rarely have significant contribution (Supplementary Figure 2B, C). Positively charged arginine residue in peptide is an interesting example: its overall contribution is stabilizing (Supplementary Figure 2A) but depending on its position within the peptide it shows effects that range from stabilizing to destabilizing (Supplementary Figure 2B). The same holds true for several other amino acids.

Overall, the largest destabilizing factor in binding across the 104 top protein-peptide models is the inability of protein to stabilize the N-terminal positively charged amino group of the peptide. However, the negative Δ*G*_bind_ values for almost all systems (101 out of 104; Figure 2A, B) show that this factor is insufficient to destabilize the overall binding of the peptide to the predicted part of the protein.

### Peptide BIP142_3 (LAEDTFGEIS) binds to CRYD protein

The 10-mer peptide fragment LAEDTFGEIS from BIP142_3/OSIP134_3 can be translated from BcTAR142/PQTAR134, expressed under stress conditions involving either *B. cinerea* or Paraquat. The above TAR is expressed solely under stress conditions and shows no expression under mock treatments (Figure 3A). We split the entire 41 AA long peptide sequence of BIP142_3/OSIP134_3 into 10-mer fragments using a sliding window and scanned against all *A. thaliana* PDB structures. High confidence peptide bindings with *p*-value less than 0.001 were retained (Figure 3B). We picked the protein-peptide model LAEDTFGEIS-2VTB(D) because the mentioned 10-mer shows sequence similarity to cryptochrome DASH (CRYD) protein (chain D in 2VTB PDB structure), hence might act as a peptide mimic or affect the activity of the protein by binding to one of its pockets. Members of the cryptochrome DASH subclade are involved in the DNA repair of cyclobutane pyrimidine dimers in single stranded DNA (Selby & Sancar, 2006). Cryptochromes in general are photolyase-like flavoproteins that mediate blue-light regulation of gene expression and photomorphogenic responses, including abiotic stress responses in *Arabidopsis*, as well as in all kingdoms of life (Yu et al., 2010).

For the LAEDTFGEIS-2VTB(D) model, binding sites were predicted using PepSite2, and a coarse model of the interacting residues between peptide and protein was built. The peptide bound to *A. thaliana* CRYD shows one of the strongest bindings among the 104 top models, with Δ*G*_bind_ value of −48.02 kcal mol^−1^. Several residues in this complex have large contribution to this binding (represented as sticks at Figure 3C), with all except N-terminal leucine of the peptide contributing in a stabilizing manner. Protein and peptide are bound via various types of interactions: salt bridges (Arg 436 and Asp 4; Arg 487, Arg 490 and Ser 10), stacking interactions (Trp 365 and Phe 6) and hydrogen bonds (Trp 365 and Thr 5). Destabilizing effect of N-terminal peptide residue, found in multiple other systems as well (Supplementary Figure 2B), is likely the consequence of the lack of a negatively charged residue at the corresponding position in the protein, which would make favourable interactions with N-terminal amine group.

Two cofactors can bind to CRYD: flavin adenine dinucleotide (FAD) and 5,10-methenyltetrahydrofolate (MTHF), out of which the first one is necessary for catalytic activity. According to UniProt database, amino acids Arg 436 and Asp 485, which coincide with LAEDTFGEIS binding site, are involved in ATP binding. If the peptide indeed binds CRYD in a way predicted in this study, it could block FAD binding or even bind simultaneously with it, therefore having an effect on the activity of this enzyme, and consequently on the aforementioned type of DNA repair (Figure 3D).

## Discussion

Steroids, peptides and other small bioactive compounds mainly regulate cellular communication in eukaryotes, including plants. Over the last decade, an increasing number of secreted peptides have been shown to influence a variety of developmental processes in plants, such as meristem size, root growth, stomatal differentiation, and organ abscission (Butenko & Aalen, 2012). sORFs that might encode peptides have been overlooked in gene prediction programs owing to their small size. Moreover, there exist only a handful of publicly available T-DNA insertion collections of peptide encoding genes (Butenko *et al*, 2014; Lease & Walker, 2006). In this study, we predict that a large number of SIPs in *A. thaliana* may exert their function through protein-peptide interactions, by binding on protein surfaces. We found 30 peptides that may bind at known ligand/cofactor binding sites on proteins. The identification of ligand/cofactor binding sites in protein structures can aid in determination of peptide ligand types and experimental validation of the function of the receptor (Glaser *et al*, 2006). A peptide may compete for the ligand-binding site on the receptor, or it may non-competitively bind to the pocket together with the ligand molecule and play a role in modulating the receptor. Additionally, we screened 15 peptides that may bind to a pocket at the interface between two monomers of a multi-chain complex. The design of peptides and peptidomimetics that mimic portions of dimeric/multimeric protein interfaces have been shown to be an useful approach for the discovery of inhibitors that bind at protein-protein interfaces (Cardinale *et al*, 2011). Currently, there is a lot of interest in drugs that can inhibit dimerization of a functionally obligate homodimeric enzyme. However, design of peptides that may disrupt protein-protein interactions is far more challenging than designing enzyme active site inhibitors, due to factors such as the large interfacial areas involved, and flat and featureless topologies that these binding surfaces may exhibit (Fletcher and Hamilton, 2006).

The characterization of protein-peptide interactions can be used to evaluate the binding affinity of the model. One of the major factors determining the binding of a peptide to a protein is the size of the pocket (Laskowski *et al*, 1996). In our study, binding analysis of the predicted protein-peptide pairs revealed that different peptides tend to bind in the same pocket of one protein. We also observed that the peptides mostly interact with proteins with the help of side chains, but this is due to the reason that we have 3.2 times more of side chain atoms than backbone atoms. However, the peptide backbone atoms participate in more unique protein-peptide interactions as compared to the side chains. This is in agreement with another finding where peptides use more H-bonds in binding to their protein partner involving the peptide backbone. In the *PeptiDB* dataset comprising of 103 protein-peptide complexes, 19 peptides bind as β-strands, which use far more H-bonds on average, while 18 peptides were bound as α-helices, which form less H-bonds with proteins and contain more nonpolar atoms at the interface (London *et al*, 2010).

The pepATTRACT-local docking method has advantages over other protein-peptide docking methods. First, pepATTRACT-local docking outperforms blind docking whose performance is similar with other local docking methods (Schindler *et al*, 2015). Second, this approach completes a run in about one hour for each pair, which is beneficial for large-scale prediction of protein-peptide interactions. In our study, 46 protein-peptide pairs failed to dock. The reason may be due to large conformational changes upon peptide binding onto the receptor, which still remains a huge problem while trying to accurately predict interactions (Trellet *et al*, 2013). Some failed cases reveal that the peptide is deeply buried into the protein surface. These failed pairs can be docked using other local docking methods. However, for most of protein-peptide interactions, only very small conformational changes upon peptide binding have been observed on the protein surface.

The *A. thaliana* genome may encode thousands of small proteins that could function as peptide signals and more than 600 plasma membrane-bound receptor-type proteins that could act as receptors for peptide ligands (Shiu & Bleecker, 2001). Several sORF-encoded peptides may target regulatory enzymes involved in metabolic pathways by downregulating or upregulating the activity of these key enzymes. Predicting potential protein-peptide pairs and confirmation of physical interaction between these pairs is crucial to advance our understanding of cell-to-cell communication during plant development or stress-tolerance mechanisms (Murphy *et al*, 2012). Nevertheless, there are a few protein-peptide pairs that have been quite comprehensively studied such as the CLE (CLV3/ESR; CLAVATA3/EMBRYO SURROUNDING REGION-related) family peptides, which are plant-specific peptide hormones that mediate cellular communication and are involved in meristem maintenance, vascular development and nematode feeding cell formation. Apart from these CLE peptides there may be many more peptides that remain to be identified. In general, our study aims at predicting potential protein-peptide interactions on protein surfaces which can be experimentally validated by researchers in the future.

## Materials and Methods

### Screening of peptide binding pockets on protein surfaces

We generated a peptide fragment library consisting of 23,113 k-mers ranging in size from 4 to 10 amino acids, using sequences of SIPs (Hazarika *et al*, 2017; De Coninck *et al*, 2013) and following a sliding window approach. We extracted 2,561 structures corresponding to 1,009 *A. thaliana* proteins from Protein Data Bank, PDB (www.rcsb.org) (Berman *et al*, 2000). 996 structures were retained after filtering out redundant ones. We carried out an all-vs-all screening of potential peptide binding sites on *A. thaliana* proteins using PepSite2 (Petsalaki *et al*, 2009; Trabuco *et al*, 2012), and retained motif matches with score > 60 and *p*-value <= 0.1. The reliability of the PepSite2 method is based on the measure of positive predictive value. For *p*-values below 0.003, false positive rate was reported to be 0.01, and true positive rate was 0.1 representing a positive predictive value of 89.9%.

We used the default settings of BLASTP2.2.28+ algorithm (Altschul *et al*, 1990) to screen out peptides that show sequence similarity with a protein chain, as entire or a part of a SIP may mimic a specific binding motif on the protein, or resemble a loop from a large structured protein, a disordered region in protein termini or interfaces between defined domains (London *et al*, 2010; Kilburg & Gallicchio, 2016).

### Building peptide models, docking and structure refinement

We shortlisted 576 protein-peptide pairs with *p*-value < 0.1 from the PepSite2 output in order to build atomistic models and perform docking studies, using the protein-peptide coarse-grained *ab initio* docking protocol pepATTRACT (Schindler *et al*, 2015). For each peptide, three idealized peptide conformations (extended, α-helical and polyproline) were built using the Python library PeptideBuilder (Tien *et al*, 2013). The backbone dihedral angles used to represent the three peptide conformations were α-helical (Φ= –57°, Ψ = –47°), extended (Φ = –139°, Ψ = –135°), and polyproline conformations (Φ = –78°, Ψ = 149°) (Trellet *et al*, 2013).

In the current study, the rigid body docking models were ranked by ATTRACT score, and the top-ranked 100 structures were subjected to atomistic refinement using the flexible interface refinement method iATTRACT. We used the distance restraint based local docking protocol of pepATTRACT to restrict the sampling during rigid body sampling stage and flexible refinement stage towards the PepSite2 predicted interface residues. The placement of peptide and protein was optimized during iATTRACT refinement. At this stage, the interface region of the peptide and the protein were treated as fully flexible, while simultaneously optimizing the center of mass position and orientation of the peptide.

## Molecular dynamics

### Preparation and parametrization

We used *pdb4amber* from Amber16 (Case *et al*, 2017) to make the pdb files of 104 high confidence protein-peptide complexes top models suitable for using this software package. All disulfide bonds detected by *pdb4amber* were retained in the system. Detected protein gaps were treated by addition of *N*-methyl group (NME) to the carbon of the backbone amide group of the C-terminal, and acetyl group (ACE) to the backbone nitrogen of the N-terminal amino acid, using PyMol Molecular Graphics System, Version v1.7.4.4, Schrodinger, LLC (Delano, 2002). Capping prevents the amino acids that are flanking the gap from being recognized as protein termini, and therefore charged.

Parametrization of the systems was done using *teLeap* from Amber16. Counter-charged ions (Na^+^ or Cl^-^) were added to the non-neutral systems, and each protein-peptide complex was surrounded by a rectangular box of explicit TIP3P water spanning 10 Å from the system. Force field *ff14SB* was used for parametrization of proteins and peptides, and *tip3p* for parametrization of water. Joung/Cheatham parameters were employed for monovalent ions in the chosen water type.

### Optimization

Systems were optimized in 25,000 steps divided in five cycles, using *sander* from Amber16. First 1,000 steps of each cycle were performed by steepest descent method, while conjugated gradient was used for the remaining steps. In first three cycles, the constraint was applied to 1. the entire protein, 2. heavy protein:peptide atoms, and 3. backbone atoms, using force constant 100 kcal mol^−1^ Å^−2^. Constraint on backbone atoms was reduced to 50 kcal mol^−1^ Å^−2^ in the fourth cycle, and no constraints were applied in the fifth.

### Molecular dynamics simulations

After optimization, each system was equilibrated during the initial 500 ps, using *pmemd* from Amber16 package. In the first 300 ps, the canonical NVT ensemble was simulated, with constraint applied to atoms in the protein:peptide complex using the force constant 25 kcal mol^−1^ Å^−2^. Temperature was increasing from 0 to 300 K during the first 250 ps. In the last 200 ps of equilibration, isothermal-isobaric ensemble NpT was simulated, with temperature held constant at 300 K and pressure at 1.0 bar, with no constraints applied to the system. Throughout equilibration, the SHAKE algorithm was used to apply constraints on bonds containing hydrogen atoms, and time step of 2 fs was used. The cutoff distance for non-bonded interactions was set to 15 Å, and the neighbor list was updated each 20 steps.

Production phase was done as a 4.5 ns continuation of the 500 ps long equilibration, using Gromacs 5 software (Lindahl *et al*, 2001; Hess *et al*, 2008; Van Der Spoel, 2005; Berendsen *et al*, 1995; Essmann, 1995; Hess, 2007; Miyamoto & Kollman, 1992; Bussi *et al*, 2007). Conversion from Amber to Gromacs file formats was performed with the help of ParmEd 2.7 tool (Swails *et al*). Constraint on bonds that contain hydrogen atoms was applied using LINCS algorithm, the time step was 2 fs, and the coordinates were written each picosecond. The temperature and pressure were kept at 300 K and1.0 bar using modified Berendsen thermostat for temperature, and Parrinello-Rahman barostat for pressure coupling. Particle mesh Ewald method□ was used for electrostatic interactions, the cutoff distance for non-bonded interactions was 12 Å, and the neighbor list was updated each 20 steps. Periodic boundary conditions were applied throughout equilibration and production phase.

### Analysis and binding energy calculation

The obtained trajectories were visualized by Visual Molecular Dynamics VMD program (Humphrey *et al*, 1996), and tools from Gromacs package were used to correct for periodic boundary conditions and calculate root mean square deviation (RMSD) of complexes’backbones. Matplotlib (Hunter, 2007) □ was used to visualize the results of analyses.

Molecular Mechanics energies with Generalized Born and Surface Area continuum solvation (MM/GBSA) method was used to calculate the Gibbs energy of protein:peptide binding in the 104 top docking models. The binding energy is calculated as the following average:

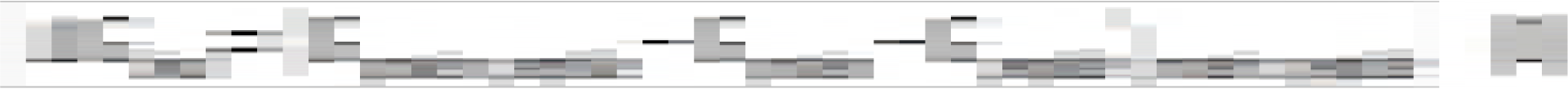

with each Gibbs energy term being the following sum:

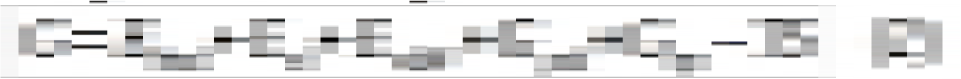

where the bonded, electrostatic and van der Waals interaction energies terms are obtained by molecular mechanics, the polar solvation term by generalized Born, the non-polar solvation term from linear relation to the solvent accessible surface area, while the entropy term is often omitted (Genheden & Ryde, 2015), as is in this study.

Amber *MMPBSA.py.MPI* was used here to calculate Δ*G*_bind_ for protein:peptide systems by MM/GBSA method, using a single trajectory of the complex. The topology files of dry complexes, as well as ligand (peptide) and receptor (protein), were prepared with Amber *ante-MMPBSA.py*. The Gibbs energy terms in equation (1) were calculated for 100 conformational snapshots from the last 2.5 ns of the production phase for each system, using salt concentration of 0.15 mol dm^−3^. During the MM/GBSA calculations, per-residue binding energy decomposition was also performed in order to get insight into contributions of specific protein and peptide residues to binding.

### Amino acids contribution to binding

The output files of the MM/GBSA per-residue energy decomposition were used to analyze the characteristics of protein:peptide binding. In each of the 104 systems, the residue with the largest contribution to binding, either in stabilizing or destabilizing manner, was detected. The threshold was then set to 40 % of its binding energy contribution value, and all residues that contributed more than the threshold in a given system were taken for the analysis, with taking into account whether amino acid belongs to protein or peptide. The number of appearances of individual amino acid was then calculated, as well as average binding contribution of different amino acids, separately for proteins and peptides, using Python.

### Characteristics of SIPs and *A. thaliana* proteins

We scanned SIPs for hydrophobicity using the grand average of hydropathy (GRAVY) number, which is a measure of the hydrophobicity/hydrophilicity of a protein based on Kyte and Doolittle equation. The hydropathy values range from −2 to +2 for most proteins, with the positively ranked proteins being more hydrophobic.

Gene ontology (GO) analysis was performed using the ClueGo Cytoscape plugin (Bindea *et al.*, 2009). Lists of 835 unique proteins from the initial screening analysis were mapped to corresponding Uniprot IDs using mappings from SIFTS database (Velankar *et al*, 2013) (www.ebi.ac.uk/pdbe/docs/sifts/index.html). The list of proteins was used to query REACTOME_Pathways and GO_BiologicalProcess ontology and the type of evidence set was All_experimental. Pathways with *p-values* ≤ 0.05 were displayed, the minimum GO tree interval was set as 3 and the maximum level was set as 8, the GO term/pathway selection was set as a threshold of 4% of genes per pathway and the kappa score was set as 0.4.

### Analysis of interactions at the protein-peptide interface

We manually inspected the top 10 models for each docked protein-peptide pair predicted by pepATTRACT using molecular visualization softwares UCSF Chimera (Pettersen *et al*, 2004) and PyMOL Molecular Graphics System. PDBeMotif, a web server for checking the PDB structure for ligands and binding sites (Gutmanas *et al*, 2014) and Catalytic Site Atlas, a database of enzyme active sites and catalytic residues on enzymes (Porter, 2004) was used for finding ligand/cofactor binding sites and enzyme active sites respectively. We analyzed if a specific peptide binding site lies at the interface of multi-chain proteins and assumed that residues on the 2 chains less than 6.0 Å apart were interacting residues. The distance between Cα atoms located in chains A and B, with coordinates A(x_1_, y_1_, z_1_) and B(x_2_, y_2_, z_2_), was calculated according to the Euclidean distance equation *D*(A,B): *√{(x*_*1*_*-x*_*2*_)^*2*^*+ (y*_*1*_*-y*_*2*_)^*2*^ *+ (z*_*1*_*-z*_*2*_)^*2*^*}.* All calculations were performed using the Biopython package from Python.

Protein-peptide bindings were characterized using BINding ANAlyzer (BINANA) (Durrant & McCammon, 2011), HBPLUS (McDonald & Thornton, 1994) and Protein-Ligand Interaction Profiler (PLIP) (Salentin *et al*, 2015) tools. BINANA was used to characterize important protein-ligand interactions such as close contacts (any receptor atom within 4.0 Å of the ligand atoms), hydrogen bonds (distance cutoff = 4.0 Å and angle cutoff <= 40°), hydrophobic contacts (ligand carbon atom within 4.0 Å of a receptor carbon atom), salt bridges and pi-pi interactions.

## Acknowledgements

RRH, NS and YS performed the analysis. RRH and VvN guided the work. This work has been supported by the KU Leuven Research Fund. NS is a doctoral fellow (1112318N) of the Research Foundation – Flanders (FWO). The computational resources and services used in this work were provided by the VSC (Flemish Supercomputer Center), funded by the Research Foundation – Flanders (FWO) and the Flemish Government – department EWI.

## Supporting Information Captions

**Supplementary Figure 1 (A)** Effect of random shuffling on the binding of peptides to pockets **(B)** Comparison of pepATTRACT-local and blind docking protocols

**Supplementary Figure 2 (A)** Average contributions to the binding energy for each amino acid type, for peptide and protein amino acids separately, and **(B)** for peptide amino acids at different locations within the peptides. **(C)** Individual data points for all amino acids, from which the averages were made, with red lines representing the average values. The represented data includes only amino acids whose binding contribution is at least 40 % of the maximal contribution value within the respective system.

**Supplementary Figure 3.**: Examples of protein-peptide models showing binding modes of characterized peptides and SIPs. The peptide binding pocket is highlighted in yellow.

**Supplementary table 1**: List of protein-peptide models where the peptide-binding pocket overlaps with known ligand binding or catalytic sites. Empty fields in the ligand column indicate only catalytic sites and no known ligand is known to bind at the respective sites.

**Supplementary table 2**: List of protein-peptide models where the peptide binding pocket lies at the subunits interface of a protein complex. For fields that are indicated as monomers in Protein stoichiometry, the other chain in the structure is either a characterized peptide or the monomers may biologically aggregate to form dimers.

